# The control of goal-directed actions by nutrient-specific appetites and rewards

**DOI:** 10.64898/2026.02.19.706921

**Authors:** Douglas J. Roy, Thomas J. Burton, Bernard W. Balleine

## Abstract

There is evidence that appetites for specific nutrients can guide foraging behaviour and aid in dietary regulation through associative learning processes that link stimuli to nutrient-specific outcomes. However, most, if not all, examples of such behaviour can be interpreted as being stimulus-bound habits, i.e., reflexive responses induced by environmental stimuli. The control of identified goal-directed actions by nutrient-specific appetites has not been directly assessed. To address this question, we trained rats to press a lever for a high protein reward (whey protein shake) and another lever for a high carbohydrate reward (polycose solution). They were then tested under extinction conditions in which both levers were available following the extended exposure to meals that were high in protein or carbohydrate. When otherwise food-deprived rats had been selectively satiated on protein immediately prior to test, they pressed more on the lever they learned had produced polycose, whereas they pressed the lever they learned had produced whey protein more if they had instead been satiated on carbohydrate. Crucially, the same pattern emerged whether the satiety manipulation was achieved using the same nutrient sources that rats had earned during training (i.e., whey or polycose) or with foods high in the relevant nutrients, indicating that these behaviours were under goal-directed control and sensitive to nutritional state. These results show that actions can be motivated by the nutritional relevance of the instrumental outcome to specific appetites, a relationship that may guide natural foraging decisions.

## Introduction

The motivational determinants of behaviour are fundamental to its explanation. Just as the legal system distinguishes between homicide and manslaughter based on intent, behavioural sciences must sometimes distinguish between “actions” and “reactions”. In this context, actions are goal-directed behaviours deployed in anticipation of a desired outcome, whereas reactions (habits or reflexes) are stimulus-bound responses provoked without the animal needing to consider their consequences. In other words, while actions are motivated by the expected value of a reward, stimulus-bound responses are driven by reinforcement history. Although these kinds of action are difficult to distinguish by observation alone, evidence suggests they are mediated by distinct learning, motivational and neural processes and are often affected differently by environmental feedback (Balleine, 2018; 2019). This makes discriminating between these forms of action important for predicting how an animal will adjust when incentive structures change and which sort of factors are likely to facilitate or impede adaptive responding.

Natural foraging situations provide particular challenges in this regard because, although a wide variety of animals are adept at organizing their behaviour to meet specific nutritional targets (Simpson & Raubenheimer, 2012; Raubenheimer & Simpson, 2020), how they achieve this remains unclear. It is possible, for example, that adaptive behaviours of this kind are merely reactions released by environmental stimuli with the threshold at which such reactions are evoked by certain stimuli modulated by nutritional state. For example, a protein-hungry animal might approach a specific food source because its associated sensory properties trigger reflexive search (e.g., Mayntz et al., 2005), whether due to a process of Pavlovian association or reinforcement of a sensory-motor association. Alternatively, animals may perform a specific action to gain access to a nutrient source because they have encoded the relationship between the action and that outcome and currently value that outcome more than alternatives.

Discriminating between these stimulus- and goal-directed explanations of specific actions has, in the past, been accomplished using post-training outcome devaluation tests to establish the degree to which an action is controlled by its relationship to and the value of a specific goal (Dickinson & Balleine, 1994; Balleine & Dickinson, 1998; Pierce-Messick & Corbit, 2024; Pierce-Messick et al., 2025). The logic of these tests is straightforward: having trained a hungry rat to perform two different actions for different food outcomes, say pressing one lever for grain pellets and another lever for sucrose solution, one or other outcome is devalued before a choice test is conducted on the two levers in the absence of the outcomes. Two methods have primarily been used to change outcome value: either the outcome is paired with illness, or it is pre-fed to induce an outcome-specific satiety effect. If, in training, the rats encoded the relationship between the actions and their outcomes and are able to integrate that information with the current value of the two outcomes then they should immediately choose to respond less on the devalued than the non-devalued lever on test, a result that has been observed many times (see Balleine, 2019 for review). Using this test, the question at issue here was whether the performance of actions associated with specific nutrient outcomes will adjust appropriately when the appetite for those nutrients has been altered. More specifically, we sought to determine if appetites for protein and carbohydrate interact with the nutritional aspects of rewards to guide goal-directed instrumental actions.

To address this issue, food deprived rats were trained to press two levers to earn two distinct rewards: one high in protein (whey) and the other high in carbohydrate (polycose). We then utilized a cross-macronutrient devaluation design using two different devaluation paradigms. First, to validate the use of these high protein versus high carbohydrate rewards, we used a traditional satiety-based outcome devaluation design, pre-feeding the rats on the one or other of the outcomes earned in training – Experiment 1A. We then repeated this test but used different protein or carbohydrate rich foods to those earned on the levers to manipulate appetite; i.e., egg or steak for selectively satiating protein appetite; cranberries or cupcakes to satiate carbohydrate appetite) – Experiments 1B and 2. By pre-feeding rats with a meal matching the macronutrient profile of just one reward but differing in its sensory properties, we hoped to assess whether the nutritional state of the rats would be sufficiently altered to change the expected values of rewards and whether their choice behaviour would be re-calibrated accordingly. Specifically, we hypothesized that, if action-oriented foraging behaviours can be based on nutrient-specific appetites, then in a choice devaluation test: (1) pre-feeding protein should selectively reduce responding on the protein-associated lever, (2) pre-feeding carbohydrate should selectively reduce responding on the carbohydrate-associated lever, and (3) that these effects will emerge whether pre-fed one of the nutrient outcomes earned in training or a sensorily distinct nutrient outcome. These results will establish that, in instrumental conditioning, nutrient-specific appetites can determine the value of nutrient-specific rewards and that choice between actions earning these rewards can be goal-directed.

## Experiment 1

### Methods

#### Subjects – Experiments 1A and 1B

The same subjects were used in Experiments 1A and 1B. Subjects were 8 male and 8 female outbred Long-Evans (250-350 g prior to start of experiment) obtained from the University of New South Wales Animal Services (Randwick, NSW). For all experiments in this paper, rats were housed in transparent plastic boxes in groups of 4 (males and females in separate boxes) in a climate-controlled colony room and maintained on a 12 h light/dark cycle (lights on between 07:00 and 19:00). All experimental stages occurred during the light portion of the day. Water and standard lab chow were continuously available prior to the start of the experiment. All experimental procedures were approved by the Animal Care and Ethics Committee at the University of New South Wales and are in accordance with the guidelines set out by the American Psychological Association for the treatment of animals in research.

Three days before the start of behavioural procedures, all rats were handled, weighed daily, and placed on a food deprivation schedule. This consisted of giving rats *ad libitum* access to water bottles in the home cage but no lab chow. Within an hour after each training or test session, rats were given combined access to lab chow and water for 1 hour in their home cages before the deprivation conditions were reimposed. Their weights were monitored daily to ensure each remained above 85% of their pre-procedure body weight throughout.

#### Apparatus – Experiments 1A and 1B

The same apparatus was used in Experiments 1A and 1B. For pre-feeding and consumption measurements in this and Experiment 1B, unless otherwise stated, the apparatus consisted of 16 feeding boxes (clear Plexiglas boxes with stainless steel lids) placed in a quiet, dark room. Foods were presented to rats in these feeding chambers in small ceramic ramekins; drinks presented in these chambers by rodent drink bottles (each made from solid plastic with stainless steel nipples). For photographic examples of materials and apparatus used in these experiments, see Supplementary Figure S1.

For all instrumental procedures, the apparatus consisted of 16 MED Associates operant chambers, each enclosed in a sound- and light-attenuating shell. The interior floor area of each chamber was 30 x 25 cm with a height of 30 cm. Each chamber was equipped with two pumps that were fitted with syringes and delivered either 0.2 mL of protein shake or polycose solution into separate wells within a recessed magazine via polyethylene tubes. Retractable levers were used for instrumental training, with rats presented with one at a time during training sessions while the other was retracted. Both levers were presented simultaneously during tests. Each lever was 48 mm wide, 19 mm deep, and set at a height 21 mm from the floor grid (lever tension: 25 g). Levers were mounted on the front panel, one on the left and the other on the right of the pellet magazine. The magazine was mounted centrally on the front panel of the chamber and consisted of two liquid cup receptacles (cup volume is .62 cc) which received liquids delivered from the pumps. A fan located on the back wall of the shell provided constant background noise (roughly 60 dB). The chambers were equipped with a sound card that delivered 90 dB 3000 Hz clicks and 90dB white noise. The walls were clear Plexiglas and the floor was a stainless-steel grid, below which was an aluminium tray for collecting subjects’ droppings. Computers running MED Associates proprietary software (Med-PC) controlled all experimental events in these chambers and recorded lever presses and magazine entries.

The outcomes in Experiment 1A used as rewards and for devaluation were mixed from whey protein concentrate, 30 g of whey protein concentrate powder added to 100 mL water plus 1 mL of vanilla essence for flavouring, and polycose (maltodextrin), 30 g of polycose added to 100 mL water plus 1 mL of vanilla essence. These substances and concentrations were chosen because of comparability in their metabolic effects and therefore likely influences on learning and choices – see Supplementary Information (Table S1) for additional information on all of the outcomes used in these experiments.

The outcomes in Experiment 1B were the same as Experiment 1A during the training phase but differed in the devaluation phase, in which alternating days of feeding were given with a high protein meal of boiled eggs with the shell removed and with a high carbohydrate meal of dried cranberries.

### Procedure – Experiment 1A

All behavioural procedures were conducted between the hours of 14:00 and 17:00 each day. Rats were fed each day with *ad libitum* access to their standard chow diet for an hour after behavioural procedures had finished for the day before otherwise being food-restricted for the duration of the experiment, with rats being returned to the colony room in their home cages without food by delivered 17:30 each day. This means that at the start of each behavioural session, rats had been deprived of food for approximately 20 hours. Rats were given 4 days to adjust to the food-deprivation regime before behavioural procedures commenced. Unless otherwise stated, all experimental work was conducted across consecutive days.

Instrumental training procedures are summarized in Table 1. Rats were given 2 days of *ad libitum* access with the instrumental outcomes to overcome neophobic reactions in their home cages prior to 2 consecutive days of magazine training with each outcome delivered in separate sessions. Rats received a magazine training session with whey protein followed immediately by a session with polycose on the first day, then again but in the opposite order on the second day. Magazine training consisted of 20 outcomes delivered on a random time 30s schedule (on average 1 outcome delivered every 30s) with each outcome accompanied by a brief auditory stimulus, either a single click or a brief burst of white noise (100 ms) counterbalanced across subjects, to ensure that reward delivery was readily detected.

**Table 1.** The design of Experiments 1A & 1B.

|  |  |  |
| --- | --- | --- |
| **Experiment 1A** |  |  |
| **Training**<br>R1→O1; R2→O2<br>(R1 & R2 were trained on alternate days for 12 days each) | **Test Day 1**<br>Pre-fed O1: R1 vs. R2<br>(The pre-feeding outcome identity was counterbalanced. Note Tests were conducted first in extinction and subsequently with the rewards delivered) | **Test Day 2**<br>Pre-fed O2: R1 vs. R2 |
| **Experiment 1B** |  |  |
| **Training**<br>Training was conducted in Experiment 1A | **Test Day 1**<br>Pre-fed O3: R1 vs. R2<br>(The pre-feeding outcome identity was counterbalanced. Note Tests were conducted first in extinction and subsequently with the rewards delivered) | **Test Day 2**<br>Pre-fed O4: R1 vs. R2 |

**Abbreviations:** R1 & R2 = left and right lever press responses (counterbalanced); O1 and O2 = whey and polycose solutions (counterbalanced); O3 & O4 = egg and dried cranberries (counterbalanced).

Following magazine training, rats were trained to lever press for protein and polycose outcomes. Levers were trained one at a time so that, for example, the right lever was presented and could be pressed to earn polycose, while in other sessions the left lever was presented to earn protein outcomes upon presses. Lever and outcome identities were counterbalanced across subjects. Each session terminated after rats earned a total of 20 outcomes or after 20 minutes had elapsed. Rats were trained to press for whey and polycose on alternating days, such that they only had a 20-minute session training them to press a lever for whey one day, a different lever for polycose the next, and so on, until rats had received 12 training sessions with each lever. Each training session was separated by a day so that rats did not confuse or misattribute metabolic effects across types of reward.

Rats were initially trained on a continuous reinforcement schedule on which each lever press was rewarded. When an individual earned all 20 outcomes of both type for two consecutive sessions, they were escalated to a random ratio 5 schedule (RR-5, reward delivered after an average of 5 presses), again until all outcomes were earned in two consecutive sessions, and then to RR-10 for the remainder of training.

#### Outcome devaluation

After instrumental training, the outcome devaluation choice test was conducted. Rats were pre-fed with one of the instrumental outcomes before they were given the opportunity to choose between both actions, with both levers presented at the same time during test. Based on studies assessing protein and carbohydrate metabolism (Chiacchierini et al., 2025; Grove et al., 2022) pre-feeding consisted of continuous free access to one of the two solutions for 30 minutes in the feeding chambers. After pre-feeding the rats were immediately placed in the operant chambers and allowed to press both levers in a choice extinction test for 5 minutes, meaning that the levers could be pressed but no outcomes would be earned. This was important because the test sought to assess whether rats were choosing to work for an outcome based on both an expectancy of the value of that outcome given current appetite conditions, and the knowledge of which specific reward each action could be expected to produce. After the 5-minute extinction test, rats were given a further 10 mins access to both levers in which they could again earn the outcomes in order to confirm that a similar pattern of behaviour emerged under the rewarded versus non-rewarded conditions.

### Procedure – Experiment 1B

From the day following the last devaluation test, the same subjects from Experiment 1A were given a brief period of “consumption training” with foods and a two-bottle choice test with whey and polycose drinks. This involved alternating days of feeding with a high protein meal (i.e., boiled eggs with the shell removed) and days fed dried cranberries in their home cages, so that rats had a total of two days of experience with each type of meal. After each 30-minute session of ad libitum access to foods, the remnants were collected and weighed in comparison with the total amount initially offered to infer the amount eaten. Nutritional values reported on food labels (summarized in Table S2 of Supplementary Information) were then used in combination with these measures to estimate how much of the target nutrients were consumed (i.e., multiplying weight of egg eaten by 0.12 to give estimated grams of protein and cranberry by 0.83 to give estimated grams of carbohydrate eaten from meals).

Rats were then given a set of consumption choice tests individually in devaluation boxes to confirm these foods were manipulating appetites for the drinks in the way intended. This choice test consisted of each subject being offered either a hard-boiled egg or cranberries in ramekins for 30 minutes before these ramekins and food remnants were removed from the feeding box and measured. Rats were then presented with two bottles, one containing the whey protein shake and the other containing the polycose solution for a further 30 minutes. Following this two-bottle choice test, the bottles were removed and measured. Rats were then offered chow in their home cages as per the food-deprivation regime and returned to the colony room before the test was repeated the following day but with the alternative type of food used to manipulate appetite: i.e., if given cranberries on day 1 they received egg on day 2 or vice versa, with the order of pre-feeding over test days counterbalanced across subjects. Results were then combined across days to enable a within-subjects analysis. The hypothesis was that egg would selectively decrease consumption of whey protein while berries would selectively decrease consumption of polycose.

Following this, the instrumental outcome devaluation tests from 1A were repeated and analysed in the same way except, in Experiment 1B, rats were pre-fed egg instead of whey protein to induce protein satiety and cranberries instead of polycose solution to induce carbohydrate satiety. We hypothesized that, in these tests, the same pattern of results observed in Experiment 1A would emerge in Experiment 1B.

### Data analysis

Due to the strong *a priori* theoretical framework guiding the design of the Nutrient-Driven Devaluation (NDD) experiment, our analysis focused on testing highly specific, directional hypotheses regarding the interaction between Appetite and Incentive. Accordingly, following the detection of a significant two-way interaction in the repeated-measures ANOVA, the hypothesis was further explored using a Simple Effects Analysis (equivalent to a set of four planned, one-degree-of-freedom comparisons). This procedure directly tested the predicted devaluation of each outcome (Simple Effect of Appetite at each Incentive level) and the predicted reversal of preference (Simple Effect of Incentive at each Appetite level).

Consistent with standard practice for testing a limited set of planned comparisons that decompose a significant interaction, the alpha criterion for each of the four simple effects tests was maintained at *p* < .05. This approach is statistically justified as the omnibus *F-*test for the interaction controls the Family-Wise Error Rate for the entire interaction effect (Maxwell & Delaney, 2004; Keppel & Wickens, 2004). Applying a post-hoc correction, such as the Bonferroni procedure, to a limited set of theoretically derived and non-exploratory comparisons would be unnecessarily conservative, significantly increasing the risk of Type II error (false negative) and reducing the power to detect the hypothesized effects (Field, 2018).

## Results and Discussion Experiment 1A

The average lever presses per session throughout instrumental training is shown in Figure 1. There was a distinct lag in presses for protein between sessions 4 and 8, attributed to technical issues with the protein pumps. While response rates for whey lagged slightly between sessions 4 and 8 due to mechanical delivery inconsistencies, rates recovered and stabilized following equipment calibration. By the conclusion of training, cumulative response rates for both outcomes had converged pressing at a rate of, on average, about 20 times per min. Repeated measures ANOVA confirmed that average press rates did not significantly differ between levers (*F*(1, 15) = .033, *p* = .859, *η^2^_p_* = .002).

**Figure 1.**
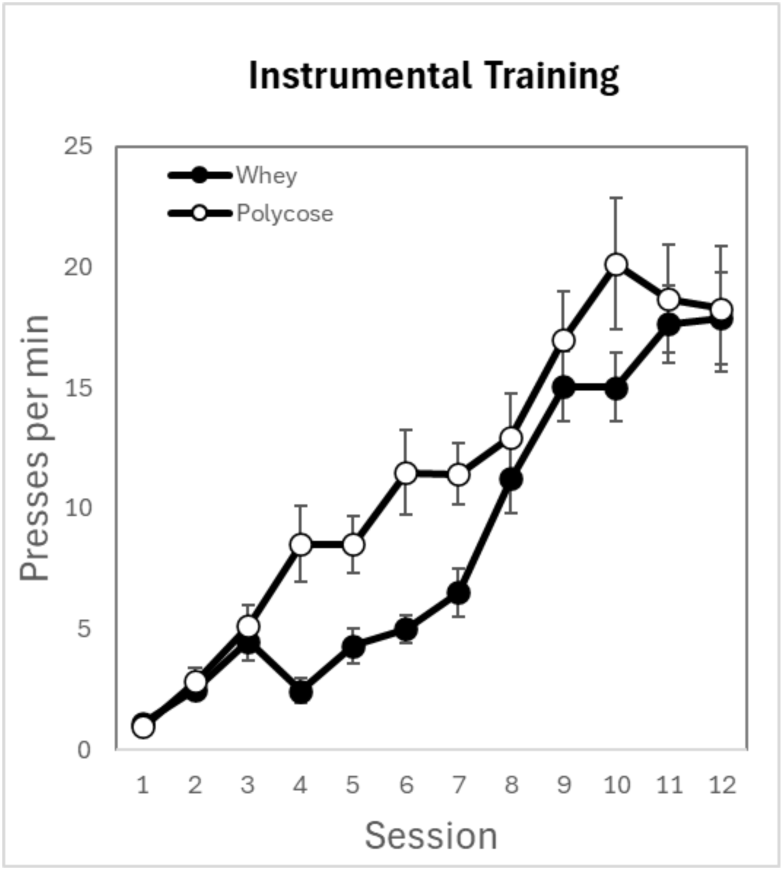
Behaviour during instrumental training. Average press rates per minute for each session during acquisition training. Error bars = ±1 SEM.

### Outcome devaluation tests

#### Extinction tests

The combined results of the outcome devaluation extinction tests are shown in Figure 2. Although rats pressed at similar rates on each lever when generally food-deprived in training, the test results clearly show rats distinguished between the types of reward and the actions that they expected were necessary to obtain them. Pre-feeding with whey appeared selectively to reduce rats’ pressing on the lever that delivered whey in training, whereas pre-feeding with polycose selectively reduced pressing on the lever associated with polycose.

**Figure 2.**
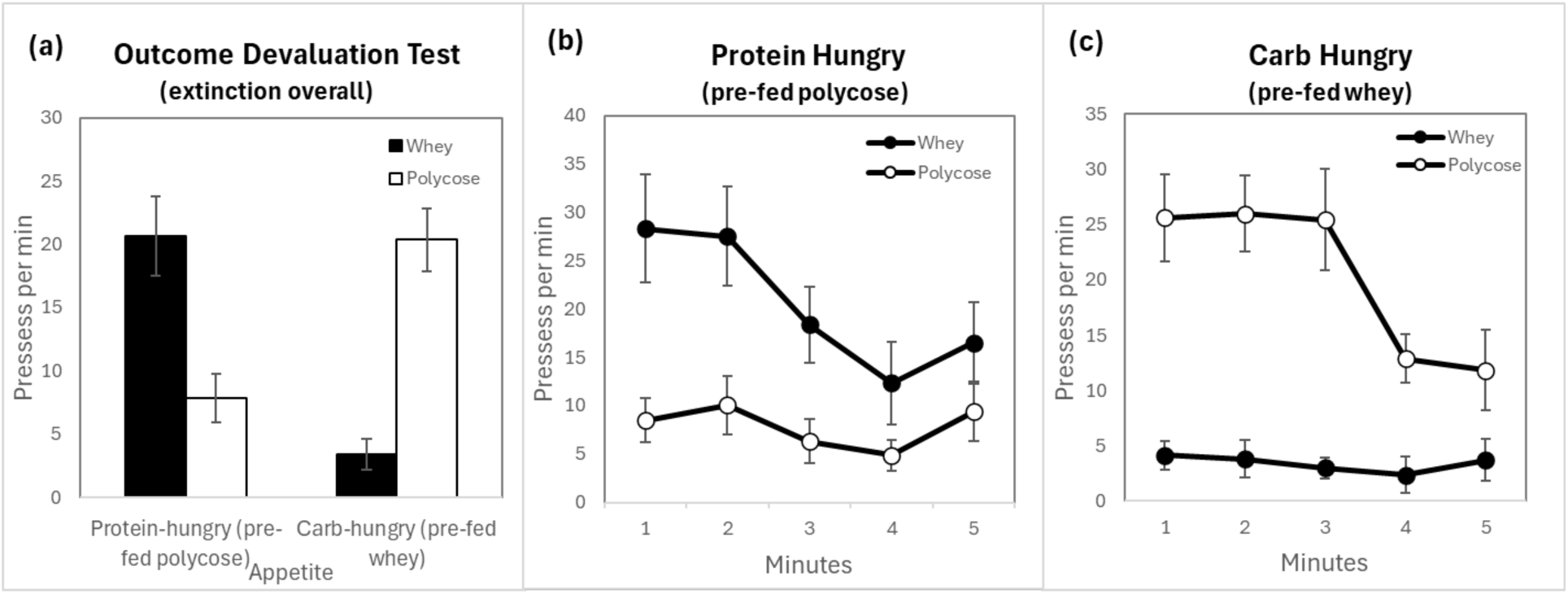
Behaviour during devaluation test under extinction conditions. (a) **M**ean press rates for the whey protein lever versus polycose lever across the two appetite conditions. Average press rates for each consecutive minute of the test are presented: **(b)** when pre-fed polycose, and (c) when pre-fed whey. Error bars = ±1 SEM

This interpretation was supported by our statistical analysis. The results of the extinction test were analysed using a 2 (Appetite: protein-hungry vs carbohydrate-hungry) × 2 (Incentive: protein vs carbohydrate lever) × 2 (Sex: male vs female) mixed repeated-measures ANOVA. There were no significant main effects of Appetite (*F*(1, 14) = 2.404, *p* = .143, *η^2^_p_* = .147), Incentive (*F*(1, 14) = 1.225, *p* = .287, *η^2^_p_* = .080, or Sex, (*F*(1, 14) = 1.113, *p* = .309, *η^2^_p_* = .074), nor any significant interactions involving Sex (all *F’*s < .926, *p’*s > .352, *η^2^_p_*′s < .062). However, the Appetite × Incentive interaction was significant, such that rats averaged more presses per minute on the lever associated with the reward when it has not been targeted for devaluation (*M* = 20.52, *SD* = 9.80) compared to when it had (*M* = 5.63, *SD* = 5.56), (*F*(1, 14) = 35.845, *p* < .001, *η^2^_p_* = .719, indicating that lever pressing was significantly influenced by the relationship between the specific appetite and outcome earned. This interaction was explored using planned simple-effects analyses (collapsing across sex). First, simple effects of Appetite were examined separately for each Incentive. Rats pressed the protein-associated lever more when protein-hungry (i.e., pre-fed carbohydrate; *M* = 20.68, *SD* = 12.61) than when protein-satiated (i.e., pre-fed protein; *M* = 3.4, *SD* = 4.78), *F*(1, 15) = 30.391, p < .001, *η^2^_p_* = .670. Conversely, rats pressed the carbohydrate-associated lever more when carbohydrate-hungry (i.e., pre-fed protein; *M* = 20.36, *SD* = 9.97) than when carbohydrate-satiated (i.e., pre-fed carbohydrate; *M* = 7.85, *SD* = 7.78), *F*(1, 15) = 22.703, *p* < .001, *η^2^_p_* = .602. Second, simple effects of Incentive were examined separately for each Appetite condition. When protein-hungry, rats pressed the protein-associated lever more than the carbohydrate-associated lever, *F*(1, 15) = 12.602, *p* = .003, *η^2^_p_* = .457. In contrast, when carbohydrate-hungry, rats pressed the carbohydrate-associated lever more than the protein-associated lever, *F*(1, 15) = 48.825, *p* < .001, *η^2^_p_* = .765.

#### Rewarded tests

Figure 3 shows the equivalent graphics for behaviour during the rewarded component of the test. The same general pattern emerged, analysed using the same methods as in the extinction test: There were no significant main effects of Appetite (*F*(1, 14) = .054, *p* = .819, *η^2^_p_* = .004), Incentive *F*(1, 14) = .579, *p* = .459, *η^2^_p_* = .040, or Sex, *F*(1, 14) = .167, *p* = .689, *η^2^_p_* = .012), nor any significant interactions involving Sex (all *F’*s < .850, *p’*s > .372, *η^2^_p_*′s < .057). The Appetite × Incentive interaction was significant, such that rats generally pressed fewer times per minute for the devalued reward (*M* = 1.61, *SD* = 1.86) than the valuable reward (*M* = 9.88, *SD* = 6.77), *F(*1, 14) =34.332, *p* < .001, *η^2^_p_* = .710, indicating that, again, lever pressing depended on the relationship between current motivational state and outcome type. Contrasts revealed an effect of appetite on presses for whey, in that rats pressed for it more times per minute when protein-hungry (*M* = 9.49, *SD* =4.58) than when carbohydrate-hungry (*M* = 1.08, *SD* = 2.17), *F*(1, 15) = 18.464, p < .001, *η^2^_p_* = .552. Rats pressed fewer times per minute for polycose when protein hungry (*M* = 2.14, *SD* =3.09) than when carbohydrate hungry (*M* = 10.26, *SD* = 6.78), *F*(1, 15) = 21.685, p < .001, *η^2^_p_* = .591. Testing when protein hungry resulted in rats showing a significant preference in press rates for the lever associated with whey over the lever associated with polycose, *F*(1, 15) =18.464, p < .001, *η^2^_p_* = .552, while testing when carbohydrate hungry had the opposite effect, *F*(1, 15) = 23.020, p < .001, *η^2^_p_* = .605.

**Figure 3.**
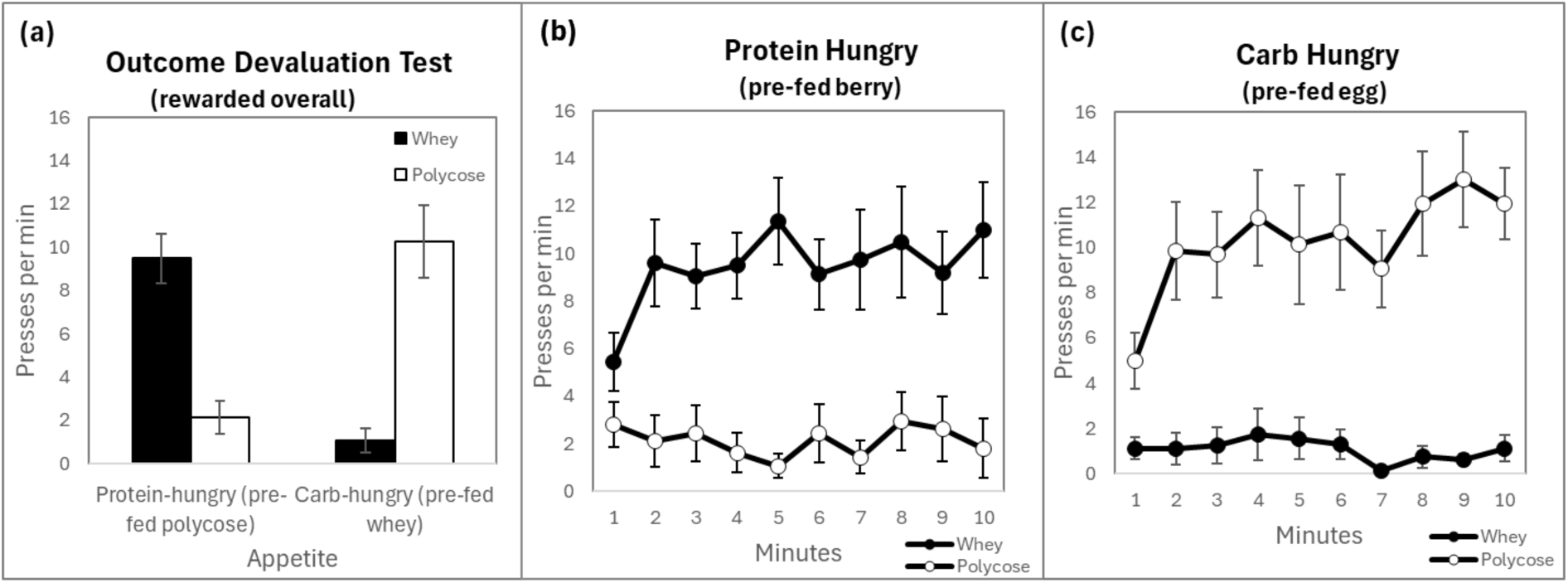
Behaviour during the devaluation test under rewarded conditions. (a) shows mean press rates for the whey protein lever versus polycose lever across the two appetite conditions. Average press rates for each consecutive minute of the test are presented for when rats were **(b)** pre-fed polycose, and **(c)** pre-fed whey. Error bars = ± 1 SEM.

### Experiment 1B

#### Consumption tests

Figure 4a shows quantities of foods consumed during consumption training (before dashed line) and during pre-feeding before choice tests (after dashed line). The results of the consumption choice test, Figure 4b, suggest these foods affected appetites in the desired directions. This interpretation was supported by a repeated measures ANOVA that analysed quantities consumed for these two types of substances across the two appetite conditions. There was no main effect of *outcome* (*F*(1, 15) =.686, *p* = .421 *η^2^_p_* = .044), however, a main effect of *appetite* was found, with rats consuming more substances during test when pre-fed cranberry (*M =* 5.56, *SD =* 2.37) than when pre-fed egg (*M =* 4.50, *SD =* 2.25), *F*(1, 15) = 5.431, *p* = .034, *η^2^_p_* = .266. There was a significant *appetite* × *outcome* interaction, suggesting that pre-feeding selectively depressed consumption of one outcome over the other, *F*(1, 15) = 12.036, *p* = .003, *η^2^_p_* = .445. Planned simple effects analysis (α=.05) clarified this pattern. Rats consumed more whey protein shake when pre-fed berries (*M =* 7.21, *SD =* 2.55) than when pre-fed egg (*M =* 3.30, *SD =* 3.05), *F*(1, 15) = 12.567, *p* = .003, *η^2^_p_* = .456, whereas they consumed less polycose when pre-fed cranberry (*M =* 3.91, *SD =* 2.19) than when pre-fed egg (*M* = 5.68, *SD* = 1.46), *F(*1, 15) = 5.750, *p* = .030, *η^2^_p_* = .277.

**Figure 4.**
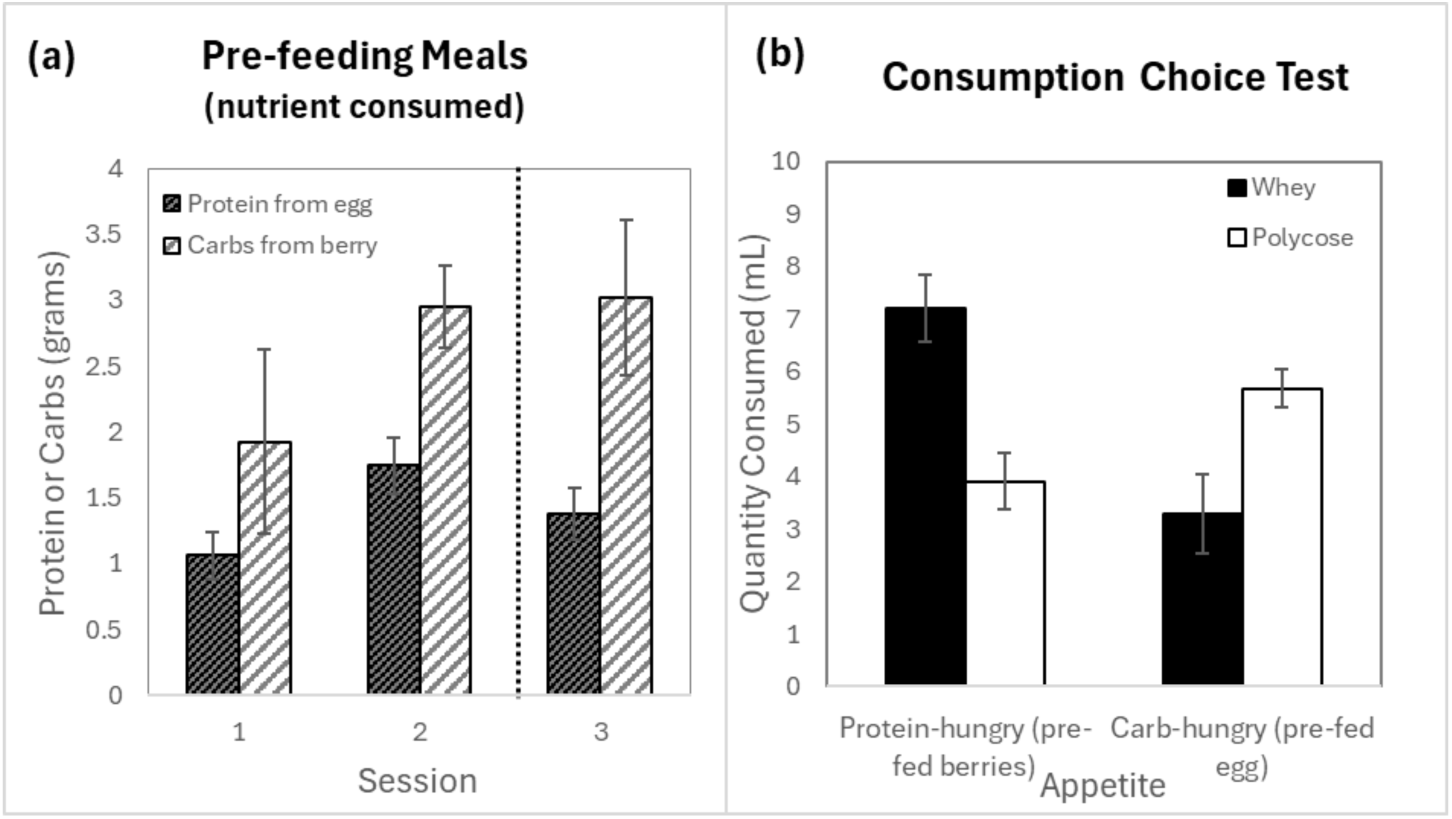
Quantities consumed during consumption training and tests using foods. **(a)** Weight of food eaten converted into the relevant nutrients they contained during consumption training and (after the dotted line) prior to a two-bottle choice consumption test. **(b)** Overall results of consumption choice test comparing how much whey vs. polycose was consumed when pre-fed a food containing the same or a different nutrient. Error bars = ± 1 SEM.

Pre-feeding with cranberries revealed a preference for whey over polycose, *F*(1, 15) =10.424, *p* = .006, *η^2^_p_* = .410, whereas pre-feeding with egg, produced a significant preference for polycose over whey (*F*(1, 15) = 6.287, *p* = .024, *η^2^_p_* = .295).

#### Extinction tests

Although not as striking, the same pattern of results in Experiment 1A were observed in Experiment 1B – see Figure 5. As can be seen in the Figure, presses on a lever were reduced when the reward with which it was associated with had been targeted for devaluation by pre-feeding: rats pressed at more on the lever trained with whey when pre-fed cranberries than when pre-fed egg and more on the lever trained with polycose when pre-fed egg than when pre-fed cranberries. This suggests that our pre-feeding with foods successfully and selectively devalued the nutritionally relevant type of reward.

**Figure 5.**
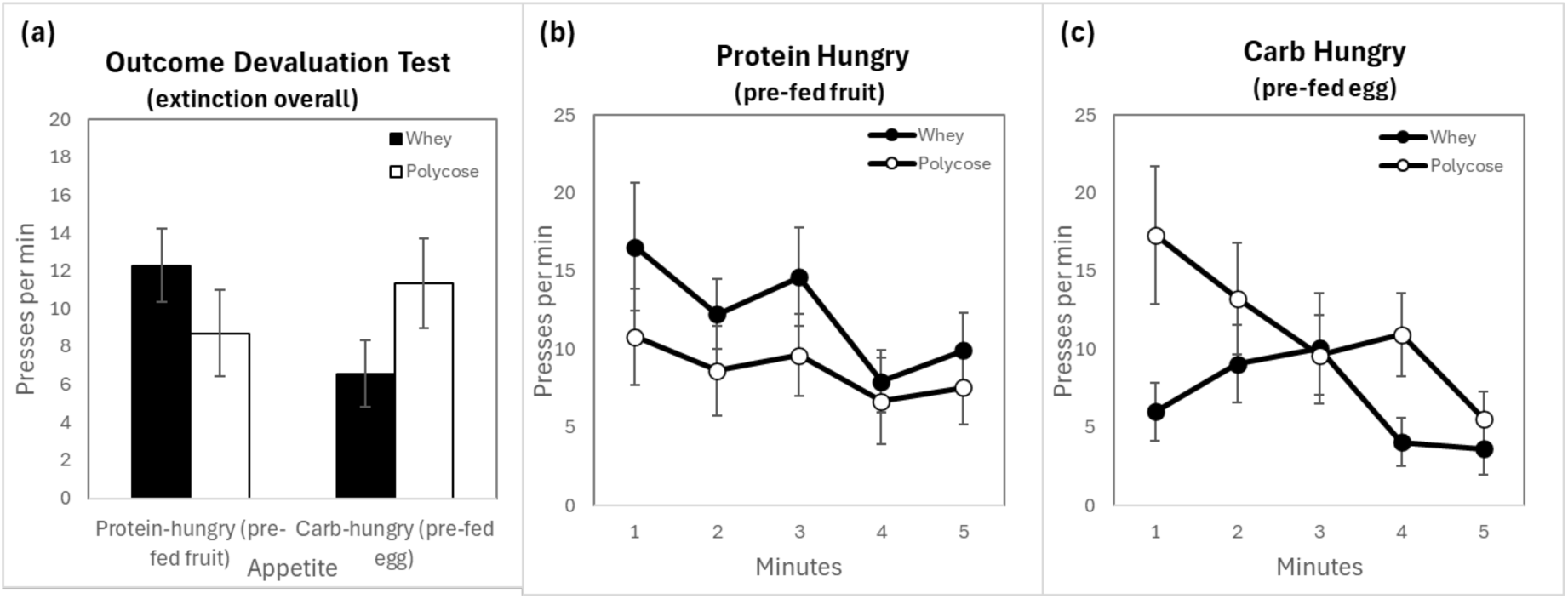
Behaviour during devaluation test under extinction conditions. **(a)** Mean press rates per minute for whey and polycose rewards under different appetite conditions. These presses are presented as averages for each consecutive minute: **(b)** when pre-fed fruit (cranberries) and **(c)** when pre-fed egg. Error bars = ± 1SEM.

Using the same mixed ANOVA as Experiment 1A, we found no significant main effects of Appetite (*F(*1, 14) =.226, *p* = .642, *η^2^_p_* = .016), Incentive (*F*(1, 14) = .098, *p* = .759, *η^2^_p_* = .007, or Sex, *F*(1, 14) = 1.059, *p* = .321, *η^2^_p_* = .070), nor any significant interactions involving Sex (all *F’*s < 2.857, *p’*s > .113, η^2’^_p_s < .170). However, we again found an Appetite × Incentive interaction, such that rats averaged more presses per minute on the lever associated with the reward when it has not been targeted for devaluation (*M* = 11.81, *SD* = 3.92) compared to when it had (*M* = 7.64, *SD* = 4.43), *F*(1, 14) = 9.754, *p* = .007, *η^2^_p_* = .411.

Despite the significant interaction, simple-effects analyses found that presses on the protein-associated lever did not differ significantly when protein-hungry (i.e., pre-fed berry; *M* = 12.28, *SD* = 7.75) vs. when protein-satiated (i.e., pre-fed egg; *M* = 6.58, *SD* = 6.97) (*F*(1, 15) =3.449, p = .083, *η^2^_p_* = .370) nor did they press the carbohydrate-associated lever significantly more when carbohydrate-hungry (i.e., pre-fed egg; *M* = 11.35, *SD* = 9.52) than when carbohydrate-satiated (i.e., pre-fed berry; *M* = 8.71, *SD* =9.01) (*F*(1, 15) = .498, *p* = .498, *η^2^_p_* = .032.). Simple effects of Incentive, examined separately for each Appetite condition found that, when protein-hungry, rats did not significantly press the protein-associated lever more than the carbohydrate-associated lever, *F*(1, 15) = 1.974, *p* = .180, *η^2^_p_* = .116; however, when carbohydrate-hungry, rats pressed the carbohydrate-associated lever significantly more than the protein-associated lever, *F*(1, 15) = 4.922, *p* = .042, *η^2^_p_* = .247.

#### Rewarded tests

Turning now to the results of the rewarded test, Figure 6, much the same pattern of behaviour emerged as under extinction conditions but with somewhat greater clarity. Mixed ANOVA revealed no significant main effects of Appetite (*F*(1, 14) =1.008, *p* = .332, *η^2^_p_* = .067), Incentive (*F*(1, 14) = .010, *p* = .922, *η^2^_p_* = .001., or Sex, *F(*1, 14) = .205, *p* = .658, *η^2^_p_* = .014), nor any significant interactions involving Sex (all *F’*s < 1.722, *p’*s > .211, η^2’^_p_s < .110). However, the Appetite × Incentive interaction was significant, *F*(1, 14) =15.283, *p* = .002, *η^2^_p_* = .522, indicating that lever pressing again depended on the match between current motivational state and outcome type.

**Figure 6.**
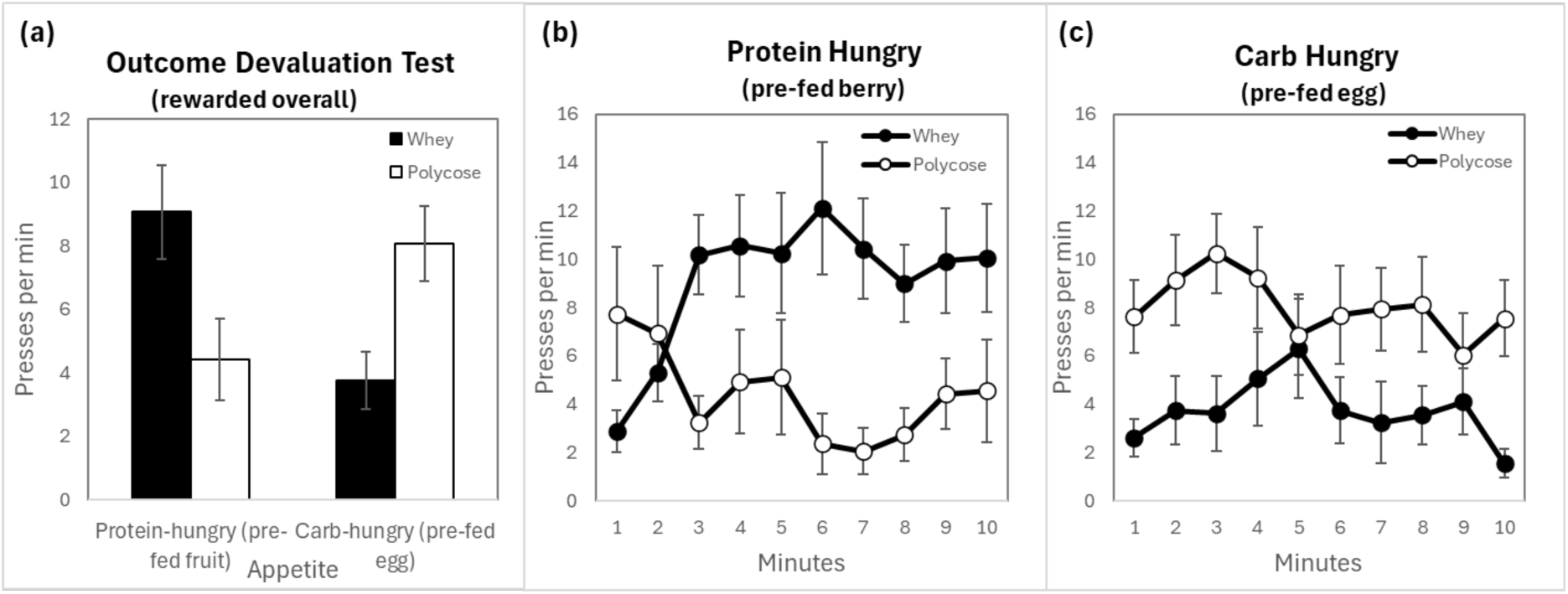
Behaviour during devaluation test under rewarded conditions using egg and cranberries to devalue whey and polycose respectively. **(a)** Mean press rates for the whey protein lever versus polycose lever across the two appetite conditions. Average press rates for each consecutive minute of the test are presented for when rats were: **(b)** pre-fed cranberries, and **(c)** pre-fed egg. Error bars = ± 1SEM.

This interaction was examined further using planned simple-effects analyses (collapsing across sex). First, simple effects of Appetite were examined separately for each Incentive. Rats pressed the protein-associated lever more when protein-hungry (i.e., pre-fed carbohydrate; *M* = 9.09, *SD* = 5.91) than when protein-satiated (i.e., pre-fed protein; *M* = 3.76, *SD* = 3.64), *F*(1, 15) = 9.438, p = .008, *η^2^_p_* = .386. Conversely, rats pressed the carbohydrate-associated lever more when carbohydrate-hungry (i.e., pre-fed protein; *M* = 8.09, *SD* =4.75) than when carbohydrate-satiated (i.e., pre-fed carbohydrate; *M* = 4.42, *SD* = 5.14), *F*(1, 15) =12.855, *p* = .003, *η^2^_p_* = .461. Second, simple effects of Incentive were examined separately for each Appetite condition. When protein-hungry, rats pressed the protein-associated lever more than the carbohydrate-associated lever, *F*(1, 15) = 4.404, *p* = .053, *η^2^_p_* = .227. Meanwhile, when carbohydrate-hungry, rats pressed the carbohydrate-associated lever more than the protein-associated lever, *F*(1, 15) = 5.331, *p* = .036, *η^2^_p_* = .262.

When considered together, the results of Experiments 1A and 1B show that, when trained to perform instrumental actions rewarded with protein vs. carbohydrate outcomes, rats can learn to associate their actions with the nutrient specific features of those outcomes and modify their performance of their actions based on the current value of those nutrients. Thus, satiation using substances high in one type of nutrient led to a preference for the reward rich in the other, as expressed in the rats’ choice both in extinction and reward choice tests.

Nevertheless, although the effects were similar in Experiments 1A and 1B, it must be admitted that the effects in the latter were not as impressive, possibly due to the same animals having experienced multiple extinction tests, which from the minute-by-minute breakdown of the extinction choice test data appear to have had a marked impact on their instrumental responding in real time. Nevertheless, it seems that manipulations of protein versus carbohydrate appetite largely explain differences in motivation to press for each type of reward, with the directionality of the effect remaining consistent across both extinction and rewarded tests. This suggests the effect is robust even if the “vigour” of the rats’ activities may have been attenuated. As the key factor involved is nutritional, we refer to this as the “nutrient-driven devaluation” (NDD) effect to distinguish it from other forms of outcome devaluation, such as those ostensibly predicated on sensory-specific satiety.

### Experiment 2

Prior experience with extinction conditions may have diminished differences in responding for rewards during the tests of Experiment 1B when they were valued versus when they were devalued, and so may have attenuated the effects in that study. We also incidentally observed that some rats ate selectively from different parts of the boiled egg: some rats selectively ate only the yolk while others ate only from the white. Egg white and yolk differ significantly in their macronutrient composition (Vliet et al., 2017). They also differ in the composition of a suite of micronutrients – many of which play yet unknown roles in metabolism (Réhault-Godbert, Guyot, & Nys, 2019) – and so it is possible that these uneven feeding patterns had uneven effects on the motivation of rats during the tests.

To address these issues, we aimed to replicate the findings from Experiment 1B but using subjects that had not already undergone extinction tests prior to the NDD test. We also used other nutritionally matched foods to target whey and polycose rewards for devaluation: For these experiments we used steak, instead of egg, to reduce protein appetite and cake, instead of cranberries, to reduce the carbohydrate appetite (see methods).

Finally, we modified the whey protein shake in two ways: First we included whey protein isolate to make it easier to mix at higher concentrations in water. Second. we also modified the source of whey protein, using commercially sweetened (saccharine) vanilla whey protein with which the whey isolate was mixed. By combining 15 grams of vanilla whey protein concentrate with 15 grams of unflavoured whey protein isolate in 100 mL of water, the whey protein shake in this experiment was estimated to be at a protein concentration of 19.4 %.

## Methods

### Subjects and Apparatus

The subjects were 16 outbred experimentally naïve Long-Evans (250-350 g prior to start of experiment – 8 male and 8 female). They were placed on food restriction a week before the start of the experiment (i.e., fed ad libitum chow in the home cage for an hour daily with food otherwise removed) to acclimate to the food-restriction regime, before a period of consumption training, followed by instrumental training and, finally, devaluation tests. The apparatus, food-deprivation, and testing regimes were the same as in Experiments 1 except where noted below (particularly regarding consumption training and the materials used for pre-feeding in the devaluation tests).

### Protein and carbohydrate outcomes for training and pre-feeding

To address mechanical delivery issues and better equate the sensory profile (sweetness) of the rewards, the whey protein shake was modified to a 19.4% concentration using a 1:1 blend of vanilla-flavoured whey concentrate and unflavoured isolate. This ensured a reliable viscosity for the pump systems while matching the high sweetness of the polycose solution (sweetened with 1% vanilla essence). Satiety was manipulated using lightly cooked beef steak (for protein) and commercially available cupcakes (for carbohydrate), providing a robust sensory departure from the liquid instrumental rewards (see Supplementary Information for more information).

### Consumption Training

The design of Experiment 2 is presented in Table 2. Rats were given 16 days of consumption training in their home cage: 8 days using protein pre-feeding and sight days with polycose pre-feeding in alternation. The eight days of protein consumption experience used lean strips of lightly cooked beef steak (see Supplementary Information for further details) presented to the rats in ramekins for 30 min. Immediately after each presentation they were given continuous free access to a bottle of either the whey protein or polycose (in counterbalanced order) for a further 30 mins. Half an hour after the end of consumption training each day, the rats were given access to chow for an hour before being returned without food to the colony room overnight. Each of these days of protein pre-feeding was alternated with eight days that were the same except that the rats were pre-fed with carbohydrate rich cupcakes, again presented in ramekins, to target carbohydrate appetite instead of protein. After each of these meals, the remnants were collected and weighed in comparison with the total amount initially offered to infer the amount eaten. Nutritional values reported on food labels were then used in combination with these measures to estimate how much of the target nutrients were consumed (i.e., multiplying weight of steak eaten by .22 to give estimated grams of protein and cake by .55 to give estimated grams of carbohydrate eaten from meals). Quantities of polyose and whey consumed were recorded to indicate whether the rats were learning to relate them to their nutrient specific appetites. Rats were given more days of consumption training than in the previous experiments to give a greater opportunity for the nutritional relationships to develop under different appetites before they were used as rewards during instrumental training. By the end of consumption training, therefore, all rats had had equivalent exposure to both whey and polycose solutions after both steak and cake pre-feeding.

**Table 2.**
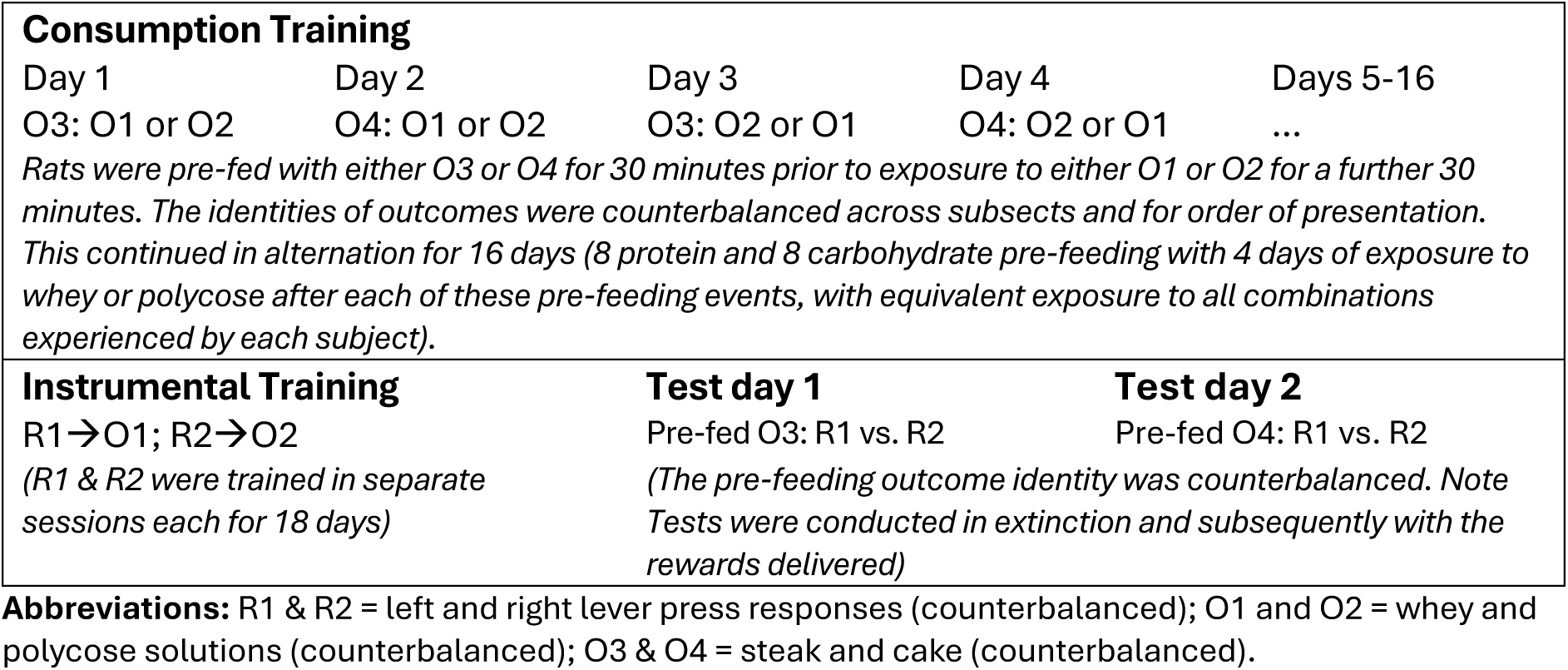
The design of Experiment 2.

### Instrumental Training and Tests

After consumption training, rats were first given magazine and lever-press training in which 45 mg grain pellets (Bioserv) were used as a common reward delivered on a continuous reinforcement schedule. This was with the hope of facilitating learning about the subsequently implemented contingencies between lever pressing and the delivery of whey and polycose outcomes. For this target instrumental training, rats were trained on a random ratio schedule of reinforcement to press for whey protein and polycose rewards as described in Experiment 1A. Each individual was trained on a continuous reinforcement schedule, followed by RR-5 and then RR-10, transitioning between schedules after successfully earning all 20 outcomes of both type for two consecutive sessions and persisting on RR-10 for the remainder of training. In this experiment, however, rats were given both training sessions on the same day. Thus, every other day, rats received a session in which they were trained to press one lever for whey protein, followed immediately by another session in which they were trained to press the other lever for polycose whereas, on the other days, the order of training sessions was reversed. The instrumental training phase lasted until all subjects successfully earned all outcomes for both rewards for four consecutive days.

Rats were then given devaluation tests as described in Experiment 1B, except using steak instead of egg and cupcakes instead of cranberries to target protein and carbohydrate appetite respectively. It was hypothesized that pre-feeding with steak would devalue whey protein as a reward relative to the polycose, whereas pre-feeding cupcakes would devalue the polycose relative to the whey protein. See Table 2 for the design.

## Results and Discussion

The quantities of foods and substances consumed during consumption training are presented in Figure 7. Regarding figure panels (a) and (b), it appeared the rats consistently consumed similar amounts of the target nutrients to the rats in Experiment 1B, despite the differing nutritional densities of egg vs steak and cranberries vs. cake (although consumption of the latter appeared more variable). By the end of training, these meals had a clear effect on the quantities of the whey and polycose outcomes consumed (panel (c)): rats consumed more whey following cake than after a steak meal, whereas they consumed more polycose following steak than after cake.

**Figure 7.**
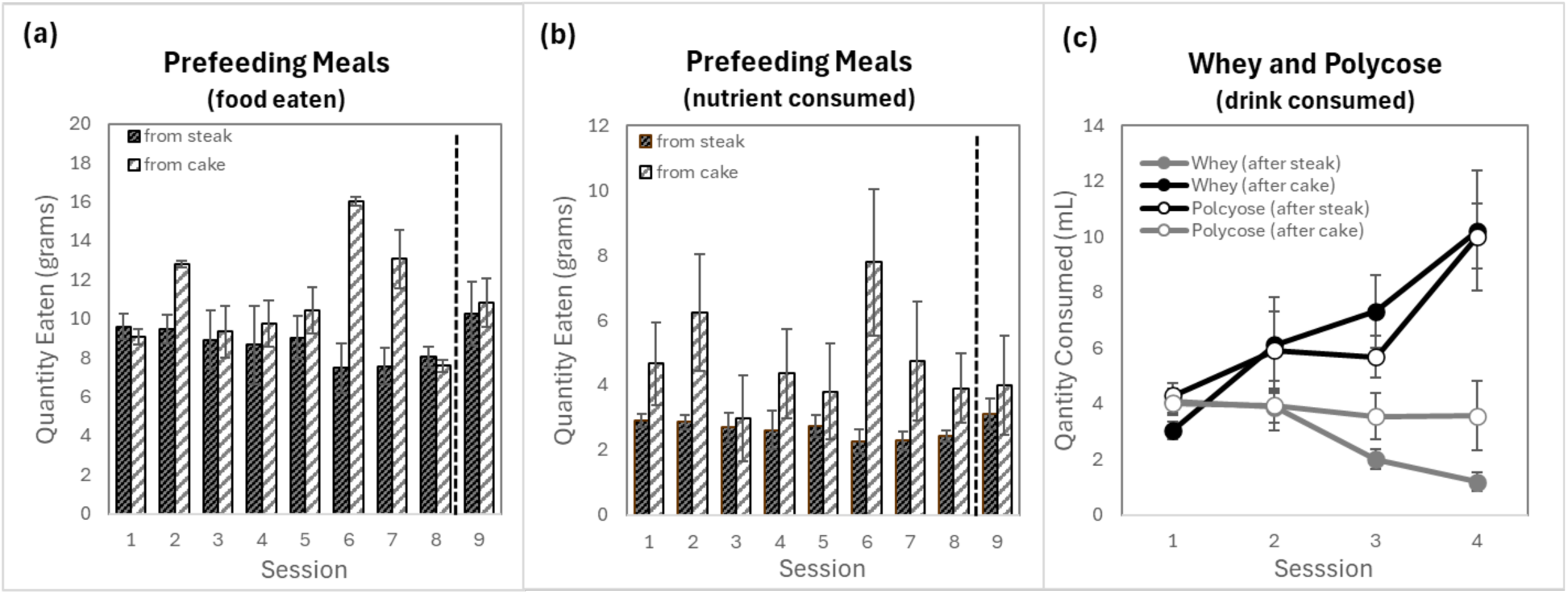
Quantities consumed of foods and drinks. **(a)** Average amount of steak or cake eaten per rat in each session during consumption training across alternating days, separated by the dotted line from quantities consumed during pre-feeding prior to the devaluation test. **(b)** Quantities of foods eaten per rat to that of their target nutrient (based on nutritional information represented on food labels). A dotted line represents the boundary between the final day of consumption training, and the quantities rats consumed as part of devaluation for the instrumental choice tests. **(c)** Quantities of drinks consumed per rat during consumption training following each type of meal (error bars = ± 1 *SEM*).

During the first session of consumption training, consumption of polycose did not differ significantly whether steak or cake was consumed during the pre-feeding (*F*(1, 3) = .173, *p* = .705, *η^2^_p_* = .055). In contrast, consumption of whey on Day 1 was significantly reduced following steak (*M* = 3.04, *SD* = .64) vs. following cake (*M* = 4.06, *SD* = .96), *F*(1, 3) = 10.494, *p* = .048, *η^2^_p_* = .778. By the end of consumption training, however, the rats consumed less whey shake (*M* = 1.19, *SD* = .66) following steak than following cake (*M* = 10.22, *SD* = 4.31) *F*(1, 3) = 23.608, *p* = .017, *η^2^_p_* = .887, and consumed less polycose following cake (*M* = 3.55, *SD* = 2.50), then following steak (*M* =10.01, *SD* = 2.33), *F*(1, 3) = 352.297, *p* < .001, *η^2^_p_* = .992.

Acquisition of the instrumental actions is presented in Figure 8. Rats appeared to learn to press for both rewards at comparable rates and a repeated measures ANOVA found no significant difference in press rates between the whey lever and polycose lever (*F*(1, 15) = 1.176, *p* = .295, *η^2^_p_* = .073).

**Figure 8.**
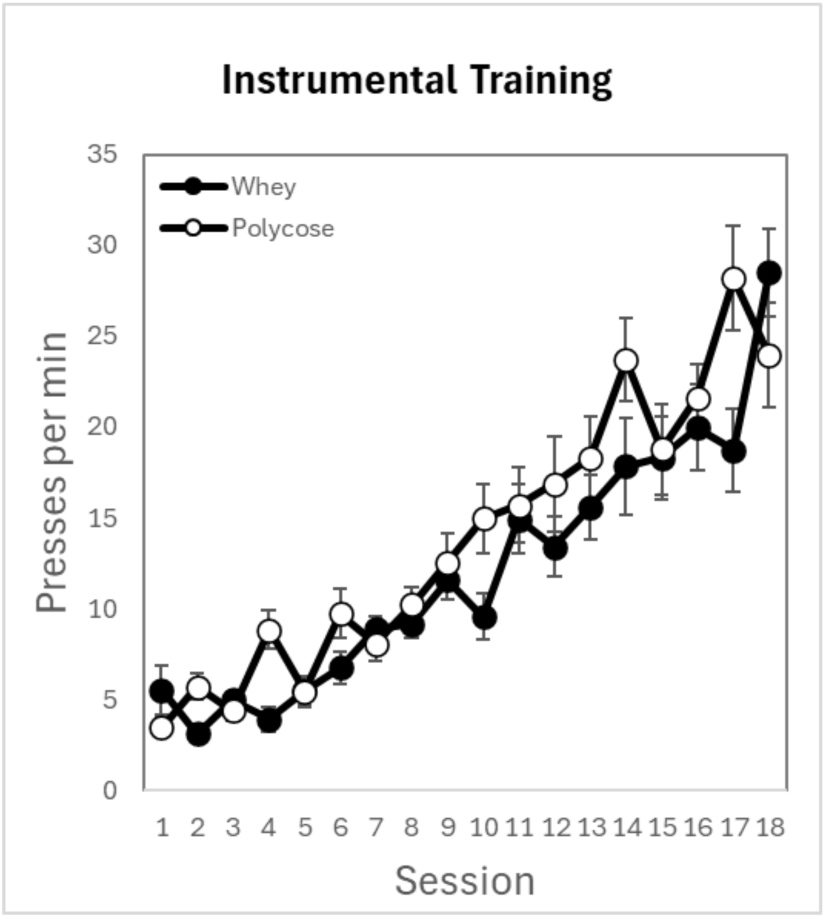
Performance on the whey and polycose levers during instrumental training. Average press rates per minute for each session during acquisition training are shown. Error bars = ±1 SEM.

Results for the choice extinction test conducted after outcome devaluation are shown in Figure 9 for each incentive across the specific appetites. On test the rats pressed the lever associated with whey more when pre-fed cake than when pre-fed steak and pressed the lever associated with polycose more when pre-fed steak than when pre-fed cake, providing a somewhat clearer replication of the results observed in Experiment 1B. The effect of the appetite manipulation appeared especially clear on the lever associated with whey.

**Figure 9.**
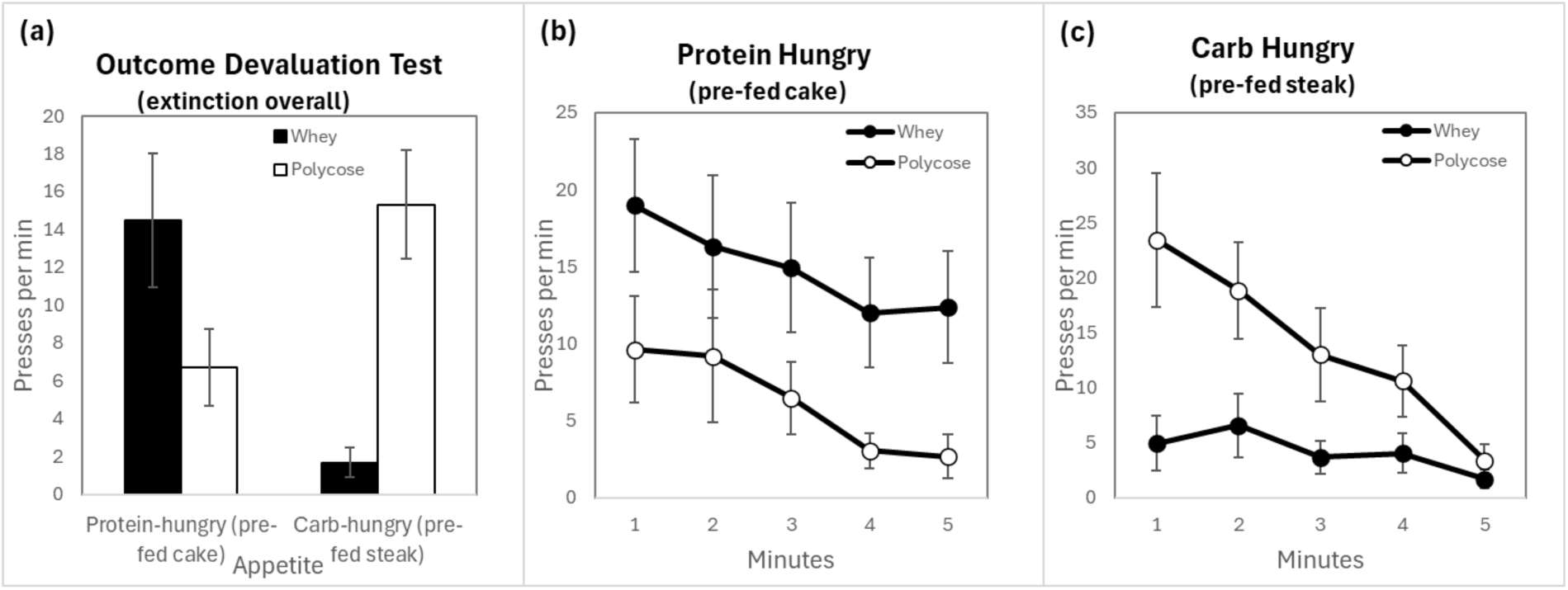
Results of the extinction devaluation test using steak and cake to devalue whey and polycose respectively. **(a)** Mean press rates per minute for whey and polycose under the different appetite conditions and with these press rates broken down into averages for each consecutive minute of the test when **(b)** pre-fed cake and so residually hungry for protein, and when **(c)** pre-fed steak, and so residually hungry for carbohydrate. Error bars = ±1 SEM.

This description was verified by a 2 (Appetite: protein-hungry vs carbohydrate-hungry) × 2 (Incentive: protein vs carbohydrate lever) × 2 (Sex: male vs female) mixed repeated-measures ANOVA. There were no significant main effects of Appetite (*F*(1, 14) = .458, *p* = .509, *η^2^_p_* = .032), Incentive (*F*(1, 14) = 1.358, *p* = .263, *η^2^_p_* = .088, or Sex, *F*(1, 14) = .440, *p* = .518, *η^2^_p_* = .030), and no any significant interactions involving Sex (all *F’*s < 1.211, *p’*s > .290, η^2’^_p_s < .080). The critical Appetite × Incentive interaction was significant: rats averaged more presses per minute on the lever associated with the reward when it has not been targeted for devaluation (*M* = 14.91, *SD* = 8.02) compared to when it had (*M* = 4.18, *SD* = 4.03), *F*(1, 14) = 19.165, *p* < .001, *η^2^_p_* = .578, indicating that lever pressing depended on the match between current motivational state and outcome type.

This interaction was further explored using planned simple-effects analyses (collapsing across sex). First, simple effects of Appetite were examined separately for each Incentive. Rats pressed the protein-associated lever more when protein-hungry (i.e., pre-fed cake; *M* = 14.49, *SD* = 14.14) than when protein-satiated (i.e., pre-fed steak; *M* = 1.68, *SD* = 3.13), *F*(1, 15) =11.238, *p* = .004, *η^2^_p_* = .428. Conversely, rats pressed the polycose-associated lever more when carbohydrate-hungry (i.e., pre-fed steak; *M* = 15.33, *SD* = 11.45) than when carbohydrate-satiated (i.e., pre-fed cake; *M* = 6.69, *SD* = 8.10), *F*(1, 15) = 4.924, *p* = .042, *η^2^_p_* = .247. Second, simple effects of Incentive were examined separately for each Appetite condition. While the direction of responding favoured the protein lever during protein-hunger, the difference did not reach statistical significance in extinction, *F*(1, 15) = 3.924, *p* = .090, *η^2^_p_* = .180, likely reflecting the high variability of initial exploratory pressing (however, this preference became robustly significant once rewards were reintroduced in the rewarded test as reported below). In contrast, when carbohydrate-hungry, rats pressed the carbohydrate-associated lever more than the protein-associated lever, *F*(1, 15) = 30.139, *p* < .001, *η^2^_p_* = .668.

As with the previous experiments, behaviour during the extinction phase of testing was similar to that observed in the rewarded phase (Figure 10). For this test the mixed repeated-measures ANOVA revealed no significant main effects of Appetite (*F*(1, 14) = 2.682, *p* = .124, *η^2^_p_* = .160), Incentive (*F(*1, 14) = .006, *p* = .938, *η^2^_p_* < .001, or Sex, *F*(1, 14) = .911, *p* = .356, *η^2^_p_* = .061), nor any significant interactions involving Sex (all *F*s < 1.364, *p*s > .262, *η^2^_p_*s < .089) but found a significant Appetite × Incentive interaction, such that rats averaged more presses per minute on the lever associated with the reward when it was not been targeted for devaluation (*M* = 7.65, *SD* = 4.46) compared to when it was (*M* =.87, *SD* =.84), *F*(1, 14) = 27.870, *p* < .001, *η^2^_p_* = .666, indicating that, again, lever pressing depended on the match between current motivational state and outcome type.

**Figure 10.**
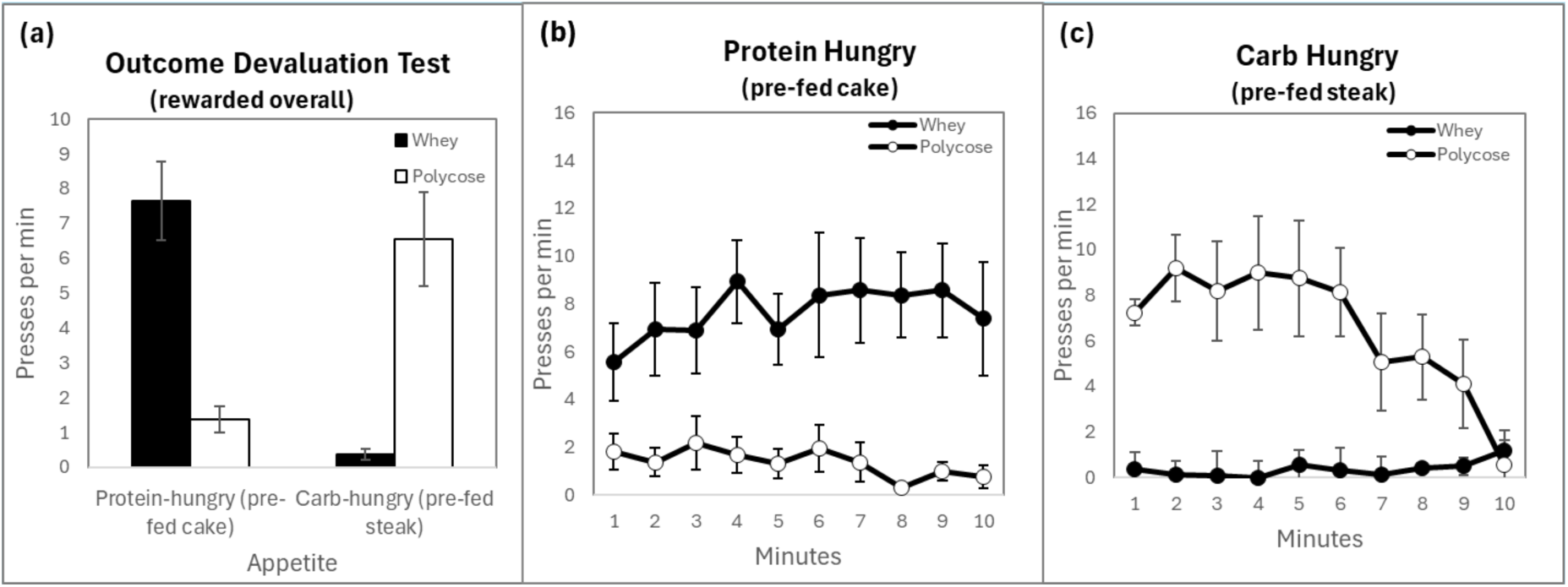
Results of rewarded devaluation test using steak and cake to devalue whey and polycose respectively. **(a)** Mean press rates per minute for whey and polycose under different appetite conditions, with these presses broken down into averages for each consecutive minute of the test when **(b)** pre-fed cake and so residually hungry for protein, and **(c)** pre-fed steak and so residually hungry for carbohydrate (Error bars = ±1 *SEM*).

Simple effects analysis found that the rats pressed the protein-associated lever more when protein-hungry (i.e., pre-fed cake; *M* =7.65, *SD* =4.46) than when protein-satiated (i.e., pre-fed steak; *M* =.37, *SD* =.64), *F(*1, 15) = 38.592, p < .001, *η^2^_p_* = .720. Conversely, rats pressed the carbohydrate-associated lever more when carbohydrate-hungry (i.e., pre-fed steak; *M* = 6.57, *SD* = 5.43) than when carbohydrate-satiated (i.e., pre-fed cake; *M* = 1.38, *SD* = 1.56), *F(*1, 15) = 12.487, *p* = .003, *η^2^_p_* = .454. Furthermore, when protein-hungry, rats pressed the protein-associated lever more than the carbohydrate-associated lever, *F*(1, 15) = 27.080, *p* < .001, *η^2^_p_* = .644 and, when carbohydrate-hungry, rats pressed the carbohydrate-associated lever more than the protein-associated lever, *F*(1, 15) = 19.818, *p* < .001, *η^2^_p_* = .569.

## General Discussion

Although it is well established that animals will perform instrumental behaviours for high carbohydrate and high protein outcomes, there has yet to be a clear demonstration that goal-directed action selection is based on the nutritive properties of the outcome. For example, protein-restricted rats have been shown to escalate the number of lever-presses to achieve a high protein reward to vastly higher “break points” than for a lower protein reward or for the same reward if returned to a normal protein diet (Chiacchierini *et al.,* 2022). Similarly, golden hamsters can learn to press a lever for protein to maintain protein intake when placed on a restricted diet, defending those levels even when the response requirement is experimentally escalated (Dibattista, 1999). Clearly, appetite for protein can invigorate activities that are instrumental in acquiring it. The present findings significantly advance this claim by demonstrating that the instrumental actions used by rats to address protein-specific requirements can be goal-directed.

Using a nutrient-driven devaluation effect (NDD), we demonstrated that the instrumental actions aimed at protein and carbohydrate acquisition are not merely “invigorated” by general hunger but can be sensitive to the specific nutritional identity of their outcomes. Because this was true even under extinction conditions – i.e., when the nutritional outcomes were not being earned as this choice was being made – rats had to base their decisions on how marginally rewarding those outcomes were expected to be if earned. The absence of immediate sensory feedback during extinction meant they also had to rely on an internal representation of the outcome, its value, and the action-outcome associations needed to achieve it (relative to alternatives). This rules out feedback from the direct detection of outcomes as driving ongoing behaviour, as occurs in consummatory choice tests.

Our design also helped to overcome limitations of the outcome devaluation paradigm that can sometimes complicate the interpretation of results. For example, null results (i.e., when response rates are equivalent for both the devalued and non-devalued outcome) can be challenging because it is difficult to discern whether such insensitivity reflects a failure of the devaluation manipulation, increased habitual responding or other factors promoting persistent responding (Meier, Staressina, & Schwabe, 2022). However, training on multiple responses with concurrent testing on two levers has been found to preclude a transition from goal-directed to habitual control; single-lever devaluation paradigms are essential to demonstrate habit formation, whereas two-lever protocols generally cannot produce or sustain habits (Kosaki & Dickinson, 2010)). Secondly, our focus was on detecting sensitivity to devaluation and, therefore, unlike investigations aiming to demonstrate an *absence* of devaluation effects due to some manipulation (e.g., silencing a suspected critical brain area), we aimed to observe the *presence* of a selective devaluation effect, emphasizing positive evidence for devaluation.

Our design also evaded the issue of sensory specific satiety (SSS) by selecting foods that differed in their sensory properties (e.g., texture, taste, appearance) from those used as instrumental rewards. The fact that pre-feeding on these foods did not significantly alter the quantities of the outcomes consumed early in the consumption training of Experiment 2, with these effects emerging over the course of this training, suggests that experience was needed for animals to recognize the relevance of each food to the value of the rewarded substances. Nevertheless, visual inspection of some of our results may suggest that pre-feeding with the same substances used in tests resulted in more marked effects on consumption and instrumental behaviour during tests. While identical-substance devaluation (Experiment 1A) showed slightly higher effect sizes than novel-food devaluation (Experiment 2), the 95% confidence intervals for these effects showed substantial overlap. The partial eta squared values for these key experiments using same substances versus different foods were 0.706 and 0.564, respectively, for the extinction tests. Both are very large, meaning that about 70 and 56 percent of the variance in lever press rates are explained by our devaluation manipulations. This suggests that the “nutritional” component of the reward value is the dominant driver of devaluation, with SSS providing a marginal additive effect. Nevertheless, the relative contribution and possible interactions between SSS and NDD are interesting and worth further investigation in future experiments.

### Implications and Future Directions

Our findings complement evidence of changes in the responsivity of reward-based neural circuitries to high protein incentives following manipulations that likely affect protein appetite. For example, rats restricted to a low protein diet show both an increased preference for high protein foods and greater activation of the nucleus accumbens, a subcortical brain area important for reward processing, following these meals than rats maintained on a normal protein diet (Tomé *et al.,* 2019). Along with the nucleus accumbens, the ventral tegmental area (VTA) forms neurocircuitry crucial for processing reward and action selection.

Chiacchierini *et al.,* (2021) conditioned rats to associate a flavour with casein protein and another with polycose and recorded neuronal activity in the VTA. They found that rats that were deprived of dietary protein developed a pronounced preference for casein over polycose. There was also a greater response in the VTA in these protein restricted rats associated with casein compared to polycose. Given our findings that protein can be a potent reward, conditionally dependent on appetite, the current results predict that other areas crucial for learning about and applying knowledge about rewards will similarly relate to protein status, such as the ventral pallidum. This area is closely connected to both the VTA and nucleus accumbens and appears to also be important for encoding the relative values of food rewards and food related stimuli (Ottenhier, Richard, & Janak, 2018; Leung & Balleine, 2013; 2015). High protein meals may similarly affect changes in dopaminergic activity and the VTA in humans. In study of obese women (Hoertel, Will, & Leidy, 2014), fasted participants were given either a high or relatively low protein meal, and changes in levels of plasma homovanillic acid in their blood was measured as an outcome (an indication of levels of dopamine production in the VTA). The researchers found that more protein consumption was associated with higher concentrations of homovanillic acid and reduced self-reported food cravings; see also Griffioen-Roose et al. (2014). Similarly suggestive is evidence that nutrient-specific satiety selectively influences eating behaviour of humans in consumption tests that are not entirely unlike those used in the consumption preparation for tests in Experiment 1B and 2 (e.g., Braden, Gwin, & Leidy, 2023; Gwin, Maki, & Leidy, 2017) as well as impacting preferences for stimuli related to specific nutrients through experience (e.g., Gibson, Wainwright, & Booth, 1995, Griffioen-Roose et al, 2012).

In the current study, NDD showed that instrumental goal-directed behaviours in rats can be influenced by the nutritional identity of incentives and that these specific appetites may plausibly operate in nature to guide foraging activities. Such an approach would conflict with prevailing views of feeding behaviour. A currently dominant notion is that there are two feeding systems. One is ‘homeostatic’, and another is ‘hedonic’, based partly on increasing evidence that carbohydrate and protein ingestion activates mesolimbic dopamine (Chaumontet, *et al.,* 2018; Journel *et al.,* 2012; Peuhkuri, Sihvola, & Korpela, 2011).

Nevertheless, these two systems are contrasted by supposing, as far back as Aristotle, that nutrient-specific appetites determine homeostatic feeding responses, whereas hedonic feeding is driven by superficial, sensory characteristics unrelated to nutrition. However, recent evidence that ‘homeostatic’ and ‘hedonic’ pathways are highly integrated functionally and anatomically (Campos, Port, & Acosta, 2022; Marinescu & Labouesse, 2024) does not support this view. A better understanding of this alternative integrative hypothesis may better align ‘hedonic’ feeding behaviour with the activity of nutrient-specific appetites, particularly in view of evidence that hedonic feedback is central to incentive learning (Dickinson & Balleine, 2009) and emerging theoretical frameworks in metabolic health, such as the recently proposed Protein Partitioning Model (Roy & Roy, 2026). Our findings provide direct behavioural evidence for such a model, demonstrating animals possess the capabilities to calibrate their efforts based on these specific nutritional requirements.

### The role of learning in nutritional regulation

In the context of this integrative hypothesis, an important question left unanswered by the above experiments is the role that learning plays in explaining these results. Procedures were designed on the assumption that rats needed familiarity with all the foods and rewards to be able to recognize how the value of these rewards varied according to appetite. For example, to know that a food reward is more rewarding when hungry than when not hungry, it is necessary rats for to have had experience consuming that reward under both high and low levels of hunger (Balleine, 1992a). In each case, we gave rats experiences with whey protein and polycose rewards under both conditions of protein hunger/carb satiety and protein satiety/carb hunger before testing whether manipulating these appetites with an acute meal or pre-feeding would selectively devalue those rewards. Whether these experiences are necessary for animals to link instrumental rewards to nutrient-specific appetites like protein-and carbohydrate-hungers, as they have been found to be for other shifts in motivation (see Dickinson & Balleine, 1994; Balleine, 2001 for reviews) is worth investigating for a variety of reasons. One is that demonstrating that such experience is necessary would further support our interpretation that our findings are explained by goal-directed processes. The consumption training data in Experiment 2 quite clearly suggest that a learning process was engaged during the course of pre-feeding coupled with protein and carbohydrate consumption. However, the degree to which this was necessary for the results of Experiment 2 was not addressed and will be an issue of interest in our future experiments.

Another residual question concerns how sources of nutrition are linked to internal states via learning. For example, it is possible that foods rapidly become linked to hunger through experience, but their relevance to nutrient-specific hungers may require much further learning to refine or specify those associations. If so, then this might help explain cases when an animal’s foraging behaviour becomes inappropriate to their nutritional needs. For instance, suppose that an animal has learned enough to know that a high carbohydrate food reward is related to hunger but has not yet had sufficient instrumental incentive learning experience to learn it is irrelevant to protein appetite. Such an individual may be inappropriately motivated by protein appetite to strive for the carbohydrate reward when it should instead be seeking out higher protein options. Factors promoting or hindering the linking of rewards to nutrient-specific hungers or other important background conditions may then plausibly play a role in explaining the development of disparities in effects related to diet as due to differences in the adroitness with which individuals can adjust their instrumental behaviours in response to challenges in their nutritional environment. For example, if the cost of one source of protein is increased, part of an adaptive response might involve shifting some efforts away from seeking that food source and instead towards alternative sources of protein. The more alternative sources an individual can correctly identify, the less vulnerable it will be to a change in the scarcity or other factors affecting the cost conditions of any one type.

Across the current experiments, our whey protein shake reward was devalued relative to polycose reward by pre-feeding on the shake itself, on egg, or on steak (three sources of similar amino acid profile and bioavailability). Although not quite as striking, we found evidence that polycose could also be selectively devalued by the polycose itself and by cranberries and cake, presumably because of their shared sugar content, even though in cranberries this is predominantly fructose and in cake it is mostly sucrose. Taken together, this gives us confidence that these devaluation effects are largely nutrient-driven. Future studies could further build this confidence by using similarly relevant foods to target other sources of protein (e.g., soy protein isolate) and carbohydrate (e.g., maltose sugar) rewards. It would be interesting to see if the phenomenon is robust to pairing earned and pre-fed foods with varying characteristics, such as rates of absorption. Studies using high protein pellets or high protein diets to probe rat behaviour almost exclusively use casein to augment protein content (e.g., Chaumontet *et al.,* 2018; Chiacchierini *et al.,* 2025). Although casein is milk-derived like whey, there are differences in the ways it affects metabolic processes leading to a much slower rise in muscle protein synthesis which might limit how well animals can relate its reward value to its current nutrient appetites (Tang *et al.,* 2009). These issues await future experiments.

## Summary and Conclusion

Outcome devaluation tests provide clear evidence that an instrumental action is goal-directed rather than driven by environmental stimuli. Using a modified form of this procedure, we showed that a high protein reward (whey) can be devalued relative to a high carbohydrate reward (polycose), using either the reward itself, or by nutritionally similar foods (and vice versa). To our knowledge, these studies comprise the first demonstration of the effect of nutrient-driven devaluation on instrumental actions earning protein versus carbohydrate rewards. It follows that at least some activities that seem driven by hunger for food in general might be open to reinterpretation as being motivated by nutrient-specific appetites. More research is needed to elucidate the learning processes involved in enabling rats to behave in this way and if similar processes operate in humans. If not only consummatory and approach responses but also goal-directed actions can be rendered sensitive to nutrient-specific appetites by learning, then this may suggest as novel target for interventions aiming to improve dietary regulation, particularly in contexts where macronutrient balance is critical such as obesity or nutrient-deficient diets.

## Acknowledgements

This research was supported by an award from the Australian Government Research Training Program to Douglas Roy and Australian Research Council grants DP200103401 to Bernard Balleine and DP240103246 to Bernard Balleine and Thomas Burton. The authors declare no conflicts of interest.

## Supplementary information

### 1 Materials and apparatus

**Figure S1.**
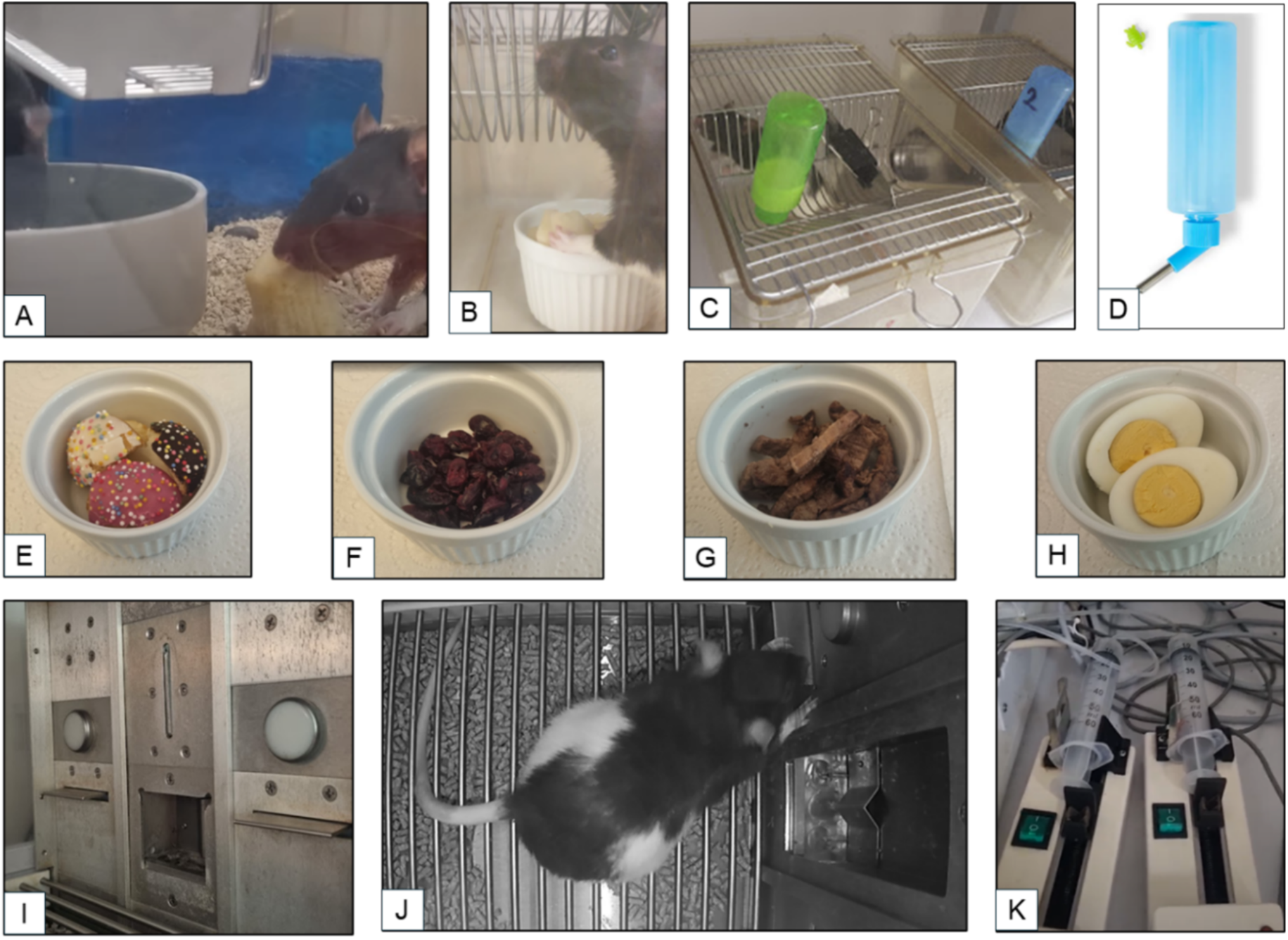
Materials and apparatus in behavioural studies. (a) Rats feeding from a large ramekin in their home cage to become familiar with cake. (b) Rat being fed individually with a ramekin in a feeding chamber. (c) Rats consuming whey protein shake from a bottle (d) stock photo of the type of bottles used for giving rats protein shakes and polycose solution drinks. (e) example of cupcakes as presented to rats in feeding chambers and home cages (f) dried cranberries in ramekin (g) lean steak in ramekin (h) boiled egg in ramekin (i) inside of the animal learning chambers showing a horizontal lever on either side of the magazine where rewards are delivered. (j) Example of a rat pressing a lever during instrumental training while the other lever is retracted (photo credit, Elise Pepin). (k) Syringes filled with drinks connected to pumps that meter out drink rewards which pass through tubing into the magazines.

### 2 Details of the nutrient-specific foods and outcomes used in these experiments

Both the whey protein and polycose solutions were designed to ensure parity in total macronutrient density and comparable metabolic salience. Details of the composition of these rewards are summarized in Table S1 and their source/procurement is listed further below.

**Table S1.**
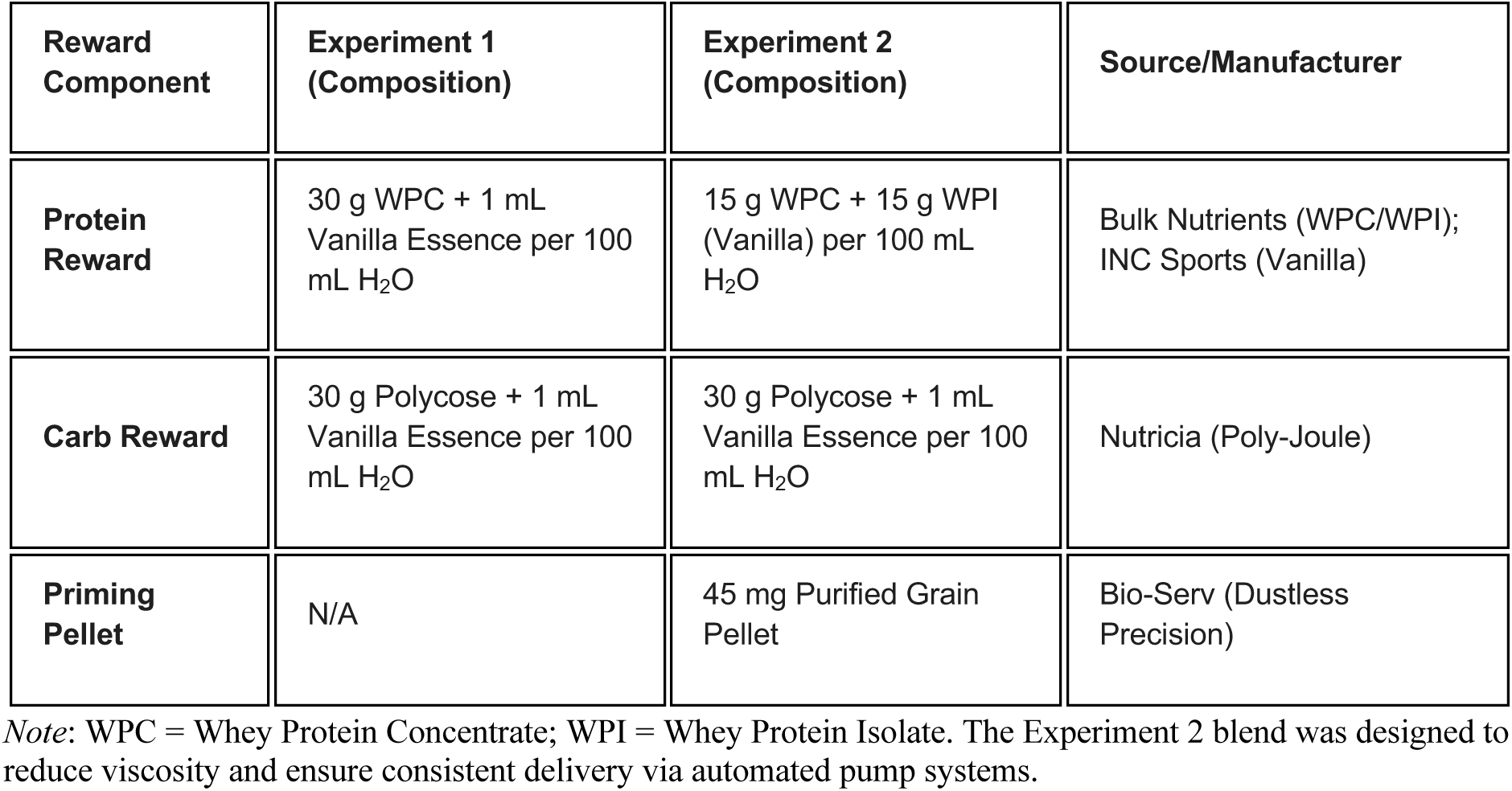

| Reward Component | Experiment 1 (Composition) | Experiment 2 (Composition) | Source/Manufacturer |
| --- | --- | --- | --- |
| **Protein Reward** | 30 g WPC + 1 mL Vanilla Essence per 100 mL H<sub>2</sub>O | 15 g WPC + 15 g WPI (Vanilla) per 100 mL H<sub>2</sub>O | Bulk Nutrients (WPC/WPI); INC Sports (Vanilla) |
| **Carb Reward** | 30 g Polycose + 1 mL Vanilla Essence per 100 mL H<sub>2</sub>O | 30 g Polycose + 1 mL Vanilla Essence per 100 mL H<sub>2</sub>O | Nutricia (Poly-Joule) |
| **Priming Pellet** | N/A | 45 mg Purified Grain Pellet | Bio-Serv (Dustless Precision) |

*Note:* WPC = Whey Protein Concentrate; WPI = Whey Protein Isolate. The Experiment 2 blend was designed to reduce viscosity and ensure consistent delivery via automated pump systems.

The concentrations used (approx. 19–22% macronutrient density) were selected to provide a potent metabolic signal while maintaining a liquid consistency suitable for instrumental delivery. High-leucine whey was chosen to maximize the rate of Muscle Protein Synthesis (MPS) (Layman, 2024). Muscle is likely an important nutrient-sensing organ for both carbohydrate and amino acids, represents a considerable proportion of the animal’s total mass, and interfaces with the brain in ways that are not yet well understood, but nonetheless seems like a strong candidate for underpinning reward signals arising from high protein rewards. In rodents, MPS typically peaks 60 minutes post-ingestion (Norton et al., 2009) (similarly for humans, see Atherton & Smith, 2012). The target dose (approximately 0.7 g protein) was achievable within a standard 20-minute session (approx. 3.5 – 4.0 mL intake), ensuring that the physiological “reward” occurred within a timeframe conducive to associative learning. An advantage of this concentration is that it would not require even aged rats to consume an enormous volume of liquid to get the required quantity of protein to trigger MPS. In similarly sized Sprague-Dawley rats, about 0.7 grams of protein from meals of whey or egg were effective at producing MPS (Norton *et al.,* 2017) which would require drinking between 3.5 - 4 mL of the whey drink used in the present study – a quantity that would likely be obtained from *ad libitum* access to the shake in rodent drink bottles, magazine training, or by earning the average of twenty rewards per session in the instrumental learning phases of the experiments and so provide what we have speculated to correspond to an important learning signal.

**Table S2.**

| Satiety Condition | Food Item (Exp) | Primary Nutrient | Protein (g/100g) | Carb (g/100g) | Fat (g/100g) | Form/ Preparation |
| --- | --- | --- | --- | --- | --- | --- |
| Protein | Hard-boiled Egg (1B) | Protein/Lipid | 12.6 | 1.1 | 10.6 | Whole, shell removed |
| Protein | Beef Steak (2) | Protein | 22.0 | 0.0 | 4.5 | Lightly pan-fried, sliced |
| Carb | Dried Cranberries (1B) | Carbohydrate | 0.1 | 82.8 | 1.4 | Raw, dried |
| Carb | Mini Cupcakes (2) | Carbohydrate | 3.8 | 55.2 | 18.0 | Sponge with icing |

*Note:* Values for Experiment 2 are derived from Woolworths product labelling. Protein-to-carbohydrate ratios were prioritized to ensure distinct motivational states. In Experiment 2, steak provided a "pure" protein signal (0% carb) to contrast with the high-sucrose load of the cupcakes.

Notably, high protein meals do not increment blood sugar levels in the same way as high glucose drinks (Franz, 1997). The metabolic effects of the high protein reward were mirror opposite to the carbohydrate reward: Polycose (maltodextrin) was chosen to induce a glycaemic excursion with a temporal profile similar to the MPS response of whey, with blood glucose typically peaking at 45–60 minutes (Lee et al., 2013) while not affecting MPS—although it may attenuate muscle protein breakdown (Børsheim *et al.,* 2004). While rewards were equated for total macronutrient concentration, we acknowledge that the thermic effect of food is significantly higher for protein (approximately 20%) than for carbohydrates (approximately 5%) (Acheson et al., 2011). Consequently, net metabolic energy may be slightly lower for the protein reward, but the temporal salience of the metabolic “pulse” was prioritized to facilitate learning.

To isolate the effects of macronutrient satiety from sensory-specific satiety, pre-feeding foods were selected for their nutritional density and sensory distinctness from the liquid rewards.

Details of these foods are summarized in Table S2. Eggs were large, free ranged eggs purchased from Woolworths supermarkets and cooked in boiling water for 3 minutes before being allowed to cool, their shells removed, then refrigerated prior to behavioural procedures. Steak was obtained in the form of Woolworths Extra Lean Beef Stir Fry. This was lightly cooked in olive oil to ensure palatability while maintaining a high protein-to-fat ratio (approximately 22% protein). Both steak and egg meals were rested outside the refrigerator and were approximately room temperature whenever presented to rats.

For carbohydrate meals, we used either dried cranberries (as in Experiment 1B), which are high in fructose (approximately 83 per cent by dry weight) and Woolworths Mini Iced Cupcakes to provide a high-sucrose load (approximately 55% carbohydrate).

### Product Source Information

- Whey protein concentrate sourced from *Bulk Nutrients.Com* with product details available at the following link https://www.bulknutrients.com.au/products/whey-protein-concentrate. Unless otherwise stated, we used the unflavoured variety.
- Polycose sources from *Nutricia* with product details at the following link https://nutricia.com.au/adult/product/poly-joule/
- Vanilla flavoured protein was *Dynamic Whey* sourced *inc. Sports Nutrition* https://incsports.com.au/products/100-dynamic-whey-vanilla/
- Pellets used in Experiment 2 were 45 mg pellets sourced from *Bioserv.* See link for product details https://www.bio-serv.com/product/DPP_RGB.html
- Details for Woolworths Dried Cranberries can be found at the following link https://www.woolworths.com.au/shop/productdetails/726369
- Woolworths Extra Learn Beef Stir Fr. See link for item and nutrition information https://www.woolworths.com.au/shop/productdetails/531015
- Woolworths Mini Iced Cupcakes (with Sprinkles). See link for item and nutrition information https://www.woolworths.com.au/shop/productdetails/360222

